# Differentiation of naïve into memory-phenotype CD8^+^ T cells does not promote the breakdown of peripheral tolerance in irradiated mice

**DOI:** 10.1101/2025.09.08.672311

**Authors:** Marine Villard, Gabriel Espinosa-Carrasco, Javier Hernandez

## Abstract

Lymphodepletion, which is currently used as adjuvant for adoptive cytotoxic T cell immunotherapy in cancer, promotes the breakdown of peripheral CD8^+^ T cell tolerance. Under lymphopenic conditions, naive T cells proliferate due to a greater availability of homeostatic cues. Proliferating CD8^+^ T cells acquire a phenotype and functionality that is similar to memory cells and are termed memory-like cells. Since memory cells are thought to have a lower activation threshold than naïve cells, it has been proposed that differentiation of potentially autoreactive CD8^+^ T cells into memory-like cells could drive the breakdown of tolerance under lymphopenic conditions. Here we studied whether lymphopenia induced proliferation and differentiation are required to overcome CD8^+^ T cell cross-tolerance in irradiated mice. Surprisingly, we found that blocking homeostatic proliferation by IL-7 neutralization did not prevent self-reactivity. CD8^+^ T cells that remained in a naïve state still became effectors upon antigen cross-presentation as efficiently as memory-like cells. Nonetheless, lymphopenia induced proliferation did enhance CD8^+^ T cell mediated self-reactivity at low T cell frequencies by increasing autoreactive T cell numbers. Thus, although homeostatic proliferation enhances CD8^+^ T cell anti-self responses, differentiation into memory-like cells is not essential for the breakdown of cross-tolerance after irradiation.

## **I**NTRODUCTION

Lymphopenia has been linked to T cell-mediated autoimmunity in a number of murine models^1,2^. Furthermore, lymphodepletion promotes anti-self-tumor antigen CTL responses in mice and cancer patients^3–5^. These observations indicate that lymphodepletion may override the mechanisms of peripheral tolerance that prevent self-reactivity. The hallmark of lymphopenia is the expansion of residual or transferred T cells. Under acute lymphopenic conditions, naïve T cells proliferate in response to an increased availability of the cytokine IL-7 and week TCR interactions with self-peptide/MHC complexes (those that induce positive selection in the thymus)^6–11^. Importantly, intermittent IL-7 signaling and week TCR engagement are two of the main factors that ensure T cell survival in normal, non-lymphopenic conditions^6,9,12^. LIP of naïve T cells is accompanied by a direct differentiation into cells that are functionally and phenotypically similar to true memory cells in the apparent absence of antigenic stimulation, termed memory-like T cells^13–16^. Despite the massive activation of CD8^+^ T cells in response to self-peptides, which could be expressed in most cells of the organism, LIP does not result in generalized autoimmunity. This is due to the existence of extrinsic and intrinsic regulatory mechanisms, TGFbeta and PTPN2, that prevent further differentiation of CD8^+^ T cells into effectors in response to these week TCR interactions^17–19^. Interestingly, TGFbeta cannot control the activation of CD8^+^ T cells in response to strong TCR signals mediated by cognate antigen^17^. Therefore, It is not surprising that self-antigen-specific T cells can induce organ-specific autoimmunity or tumor rejection under lymphopenic conditions^5,20,21^. The commonly accepted explanation for these results is that self-antigen specific memory-like CD8^+^ T cells generated upon LIP have less stringent requirements for activation and thus may escape tolerization upon self-antigen encounter^22,23^. This hypothesis found support in the fact that memory-like T cells are less prone to tolerization than naïve cells^24^. However, it remains to be proved whether LIP and differentiation into memory cells are required for the breakdown of CD8^+^ T cell tolerance.

We have previously shown that potentially autoreactive naive CD8^+^ T cells undergo deletional tolerance, even in the presence of naïve antigen-specific CD4^+^ T helper cells, upon self-antigen cross-presentation^25,26^. This was evaluated in adoptive transfer experiments of inluenza virus hemagglutinin (HA)-specific TCR transgenic Clone 4 CD8^+^ T cells and HNT CD4^+^ T cells into InsHA mice wherein HA is expressed under the control of the rat insulin promoter in the beta cells of the pancreas. Importantly, the outcome was completely different when T cells were transferred into hosts rendered lymphopenic by mild irradiation^21,27^. In these conditions, CD8^+^ T cells were able to overcome cross-tolerance and induce autoimmunity although this effect was dependent on the presence of HA-specific CD4^+^ T helper cells^21^. In our lymphopenic model, both HA-specific CD8^+^ and CD4^+^ T cells underwent extensive LIP and differentiated into memory-like cells. Memory-like CD4^+^ T cells promoted the further differentiation of memory-like CD8^+^ T cells into effector CTL in response to antigen cross-presentation and their migration to the site of antigen-expression^21^. In the absence of help, however, despite of undergoing extensive LIP and differentiation into memory-like CD8^+^ T cells they were not able to overcome tolerance in the draining lymph nodes (LN) of the pancreas^21^. In this study, in light of these previous results and given the fact that there is no direct evidence demonstrating that CD8^+^ T cell LIP and differentiation are essential for overriding tolerance, we have addressed this issue in our mouse model of lymphopenia-induced self-reactivity. We found that differentiation into memory-like CD8^+^ T cells was not required to overcome CD8^+^ T cell cross-tolerance. Differentiation of CD8^+^ T cells into effector CTL was equally efficient after LIP blockade.

## MATERIALS AND METHODS

### Mice

BALB/c mice were purchased from Charles River. InsHA^28^, Clone 4 TCR Thy1.1^+/+29^ and HNT TCR Thy1.1^+/+30^ transgenic mice lines were propagated and maintained under specific pathogen-free conditions in the Institut de Génétique Moléculaire de Montpellier animal facility. Experimental procedures were approved by the Animal Care and Use Committee Languedoc-Roussillon, CEEA-LR-12163.

### T cell isolation

Naïve CD8**^+^** T cells from Clone 4 TCR Thy1.1 transgenic mice and CD4^+^ T cells from HNT Thy1.1 were prepared from single cell LN and spleen suspensions by magnetic depletion using the T CD8^+^ and T CD4^+^ negative isolation kits (Dynabeads, Invitrogen) according to the manufacturer’s instructions. T cell purity was greater than 85%. Purified Clone 4 CD8^+^ T cells display a homogeneous naïve phenotype^21^. In some experiments, purified T cells (2×10^7^ cells/ml) were labeled with 2 µM 5- and 6-carboxy-fluorescein succinimidyl ester (CFSE) (CellTrace™ CFSE Cell Proliferation Kit, Invitrogen)^21^. Tregs from Balb/c mice were purified from negatively isolated CD4^+^ T cells by the use of the CD4^+^ CD25^+^ positive isolation kit (Miltennyi).

### Mice irradiation and adoptive transfer

BALB/c or InsHA mice were sublethaly irradiated (450 rads) utilizing a therapeutic irradiator, Varian. Mice were used for adoptive T cell transfer experiments 24 h after irradiation.

### *In vivo* treatments with antibodies and cytokines

BALB/c or InsHA mice received every other day 0.5 mg of anti-IL-7 mAb^31^ (clone M25, BioXCell) and 0.5 mg of anti-IL-7Ralpha mAb^32^ (clone A7R34, BioXCell) by i.p. injection starting at 24 h after irradiation and up to day 7. For long term experiments, mice were then injected every 4 days. Control mice received the same amount of the matched isotype control antibodies (clone MPC-11, mouse IgG2b; clone 2A3, rat IgG2a, BioXCell).

IL-7 immune complex were prepared by mixing 1.5 µg of anti-IL-7 mAb M25 and 7.5 µg of carrier free recombinant mouse IL-7 (eBiosciences) in 200 microliters of PBS per mouse per dose^33^. IC were injected i.p. 2, 4 and 6 days after adoptive transfer.

### Diabetes monitoring

Mice were monitored for diabetes by measuring blood glucose every 3 days for a maximum period of 30 days after T cell transfer with a glucometer Breeze 2 Apparatus (Bayer, France).

### Flow cytometry

Pancreas, spleen, pancreatic (pLN) and a mixture of inguinal, axillary, cervical, mandibular, popliteal and mesenteric LN were excised and processed to obtain single cell suspensions. Staining and analysis on a FACSCanto II apparatus (BDB, Mountain View, CA) were performed as previously described^21^

All mAbs and secondary reagents were purchased from BD PharMingen (San Diego CA) except anti-CD25-APC-eFluor® 780, anti-GranzymB-PE, anti -Tbet-APC and anti-Foxp3-APC mAbs (eBioscience, San Diego, CA). Donor Clone 4 CD8^+^ T cells were detected and enumerated by virtue of Thy1.1 expression with anti-CD8α-V500 and anti-Thy1.1-PerCP mAbs. To assess the phenotype, they were stained with anti-CD25-APC-eFluor® 780, anti-CD62L-APC, anti -CD122-PE, and anti-CD44-PE-Cy7 mAbs. mAbs. Intracellular Granzyme B, T-bet and Foxp3 staining was performed utilizing the Fixation and Permeabilization Kit (eBioscience) according to manufacturer’s instructions. Matching isotype labeled antibodies were used as controls for specific staining. Intracellular IFNgamma was detected with anti-IFNgamma-PE and the BD Cytofix/Cytoperm™ Fixation/Permeabilization kit after single cell suspensions were stimulated with peptide as described^21^.

### Statistical analyses

Statistical significance was determined using a Student’s *t* test with a one-tailed distribution and two-sample equal variance. For pancreas infiltration experiments significance was evaluated using the chi square test. Data were considered to be statistically different (*) for *P* <0.05.

## RESULTS

### IL-7 blockade prevents LIP and differentiation of Clone 4 CD8^+^ T cells in syngeneic hosts

As we previously reported^21^, naïve HA-specific Clone 4 CD8^+^ T cells undergo extensive LIP when transferred into sublethaly irradiated, syngeneic, antigen-free hosts (Fig. 1A). Proliferation in irradiated mice is mediated by the increased availability of homeostatic cues, including IL-7^34,35^. Thus, preventing IL-7 signaling has been shown to efficiently block LIP and differentiation into memory-like cells *in vivo*^36^. In order to block LIP, we utilized M25 anti-IL-7 and A7R34 anti-IL-7Ralpha monoclonal antibodies (mAbs). These well-characterized mAbs have been extensively utilized to block IL-7 signaling with no significant signs of T cell depletion^36–38^. Indeed, treatment of irradiated BALB/c mice every other day with 0.5 mg of each M25 and A7R34 (IL-7 blockade hereafter) prevented LIP of transferred Clone 4 CD8^+^ T cells in secondary lymphoid organs (Fig. 1A). Accordingly, total numbers of recovered donor T cells were significantly 5-fold lower in treated than in control mice that received an isotype matched mAb (Fig. 1B). Notably, IL-7 blockade also prevented LIP of HA-specific HNT CD4^+^ T cells in syngeneic irradiated mice (Sup. Fig. 1).

**Figure 1.**
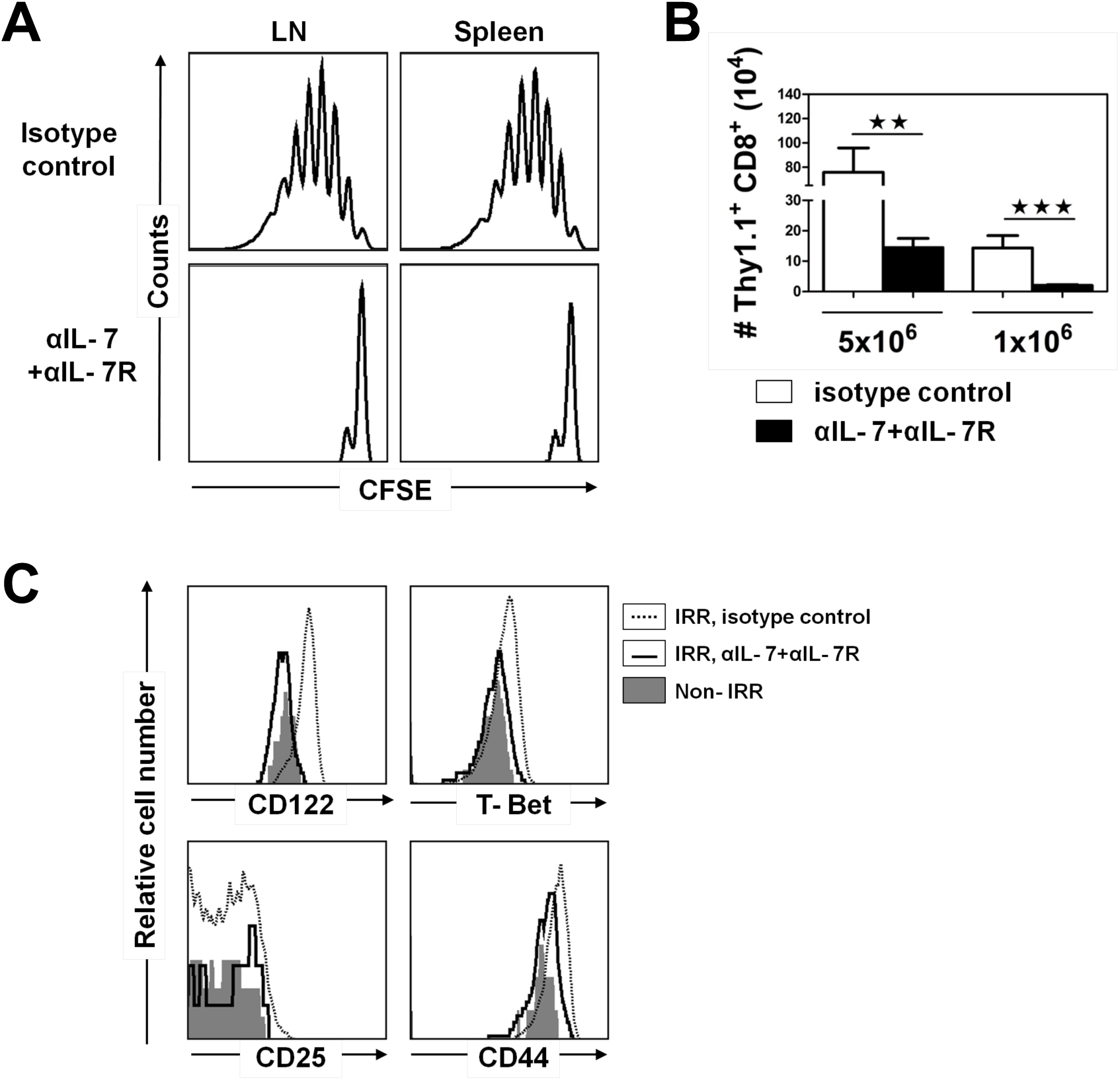
IL-7 blockade prevents LIP and differentiation of naïve Clone 4 CD8^+^ T cells into memory-like T cells. (**A)** Irradiated BALB/c mice were injected with 5×10^6^ CFSE-labeled naïve Clone 4 Thy1.1^+^ CD8^+^ T cells and 5×10^6^ HNT CD4^+^ T cells and treated with either anti-IL-7 and IL-7Ralpha or isotype control mAbs. Mice were sacrificed at day 8 after transfer and cells from the LN and spleen were analyzed by FACS. Histograms represent CFSE labeling on gated CD8^+^ Thy1.1^+^ lymphocytes. (**B)** Total numbers of CD8^+^ donor T cells in the lymphoid organs were enumerated at day 8 post-transfer following the injection of the indicated amount of Clone 4 CD8^+^ T cells and HNT CD4^+^ T cells together with isotype control (□) or with anti-IL-7 and IL-7Ralpha mAbs (▪). Results are presented as means ± SD (*n* =5-7). (**C)** CD8^+^ Thy1.1^+^ from the LN of irradiated BALB/c mice described in **A** and of adoptively transferred non-irradiated BALB/c mice were assessed for expression of the CD44, CD25, CD122 and T-bet at day 8. Irradiated treated mice are presented as solid line histograms, irradiated isotype controls are pointed lines and non-irradiated mice are shaded histograms. Data from one representative experiment of 4 are presented except for **B** where cumulative data from 2 independent experiments are presented.

Proliferating naive Clone 4 CD8^+^ T cells differentiate into cells that resemble central memory in irradiated BALB/c mice^21^. They upregulated CD122 and CD44, while they did not upregulate the effector cell marker CD25 respect to naïve cells (Fig. 1C). They also upregulated the transcription factor T-bet, as recently reported for memory-like cells^36^ (Fig. 1C). IL-7 blockade prevented the upregulation of CD122, CD44 and T-bet. Indeed, Clone 4 CD8^+^ T cells recovered from treated mice displayed a phenotype that was almost indistinguishable from naïve cells (Fig. 1C). It is important to note that the population of naïve Clone 4 CD8^+^ T cells presented a homogeneous phenotype and that no memory-phenotype T cells were detected (^21^ and Fig. 1C). Our results indicate that proliferation of HA-specific TCR transgenic T cells in irradiated hosts is fully dependent on IL-7 and that blocking IL-7 efficiently prevents Clone 4 CD8^+^ T cell LIP and differentiation into memory-like cells.

### IL-7 blockade does not prevent the breakdown of CD8^+^ cross-tolerance in irradiated InsHA mice

Cotransfer of naïve HA-specific Clone 4 CD8^+^ T cells together with HNT CD4^+^ T cells into sublethaly irradiated InsHA mice promoted the breakdown of cross-tolerance and the onset of autoimmune diabetes^21^ (Table 1). In this model, lymphopenia induced the proliferation and differentiation of both potentially autoreactive CD8^+^ as well as CD4^+^ T cells into memory-like cells. Upon antigen encounter in the draining lymph nodes of the pancreas, memory-like CD8^+^ T cells further differentiated into effector CTL with the help of HA-specific CD4^+^ T cells. Then, CTL migrated to the pancreas and mediated beta cell destruction^21^. To determine whether LIP and differentiation of Clone 4 CD8^+^ T cells into memory-like cells is required for the breakdown of CD8^+^ T cell cross-tolerance and their further differentiation into effector CTL, we targeted the IL-7 pathway in irradiated InsHA mice. Lymphopenic InsHA mice, coinjected with equal numbers (5×10^6^) of Clone 4 CD8^+^ and HNT CD4^+^ T cells, were treated with the M25 and A7R34 mAbs. Surprisingly, all treated mice still developed autoimmune diabetes (9/9) (Table 1).

**Table 1.** IL-7 blockade only reduces self-reactivity at low T cell frequencies.

| Host | Cell transfer<sup>a</sup> | Treatment<sup>b</sup> | Diabetes incidence<sup>c</sup> |  |  |
| --- | --- | --- | --- | --- | --- |
| **Irradiated InsHA** | 5x10<sup>6</sup> | isotype control | 100% | n=9 | d13±3 |
|  |  | αIL-7+ αIL-7R | 100% | n=9 | d11±1 |
|  | 10<sup>6</sup> | isotype control | 71% | n=24 | d14±4 |
|  |  | αIL-7+ αIL-7R | 38% | n=16 | d12±2 |
| **Non- irradiated InsHA** | 5x10<sup>6</sup> | isotype control | 0% | n=10 |  |

<sup>a</sup> Equal numbers of purified transgenic Clone 4 CD8<sup>+</sup> and HNT CD4<sup>+</sup> T cells were injected as indicated into irradiated or non-irradiated InsHA mice, 24h after irradiation.
<sup>b</sup> Mice were treated either with anti-IL-7 and anti-IL-7Ralpha mAbs or isotype control mAbs
<sup>c</sup> The onset of autoimmunity was evaluated by measuring blood glucose levels. Mice were followed over a 30 days period and were considered diabetic when levels were above 300 mg/dl in two consecutive measurements. The day of disease onset (d) is indicated when applicable.

CFSE profiles of donor CD8^+^ T cells showed that IL-7 blockade prevented LIP, as evidenced in non-draining LN, compared to isotype control mice. However, Clone 4 cells proliferated extensively in response to HA antigen cross-presentation and CD4 help in the pancreatic (p)LN of treated and isotype control mice (Fig. 2A). Furthermore, migration of donor CD8^+^ T cells to the pancreas was not significantly different in anti-IL-7/IL-7R treated as compared to isotype control mice (Fig. 2A and 3). These proliferation profiles were in clear contrast to those of naïve Clone 4^+^ CD8^+^ T cells in the pLN of non-irradiated InsHA mice, where they undergo deletional cross-tolerance even in the presence of naïve HNT CD4^+^ T cells^26^. Cells proliferated slowly in the pLN, did not accumulate as they proliferated and did not recirculate to other lymphoid tissues or migrated to the pancreas (Fig. 2A).

**Figure 2.**
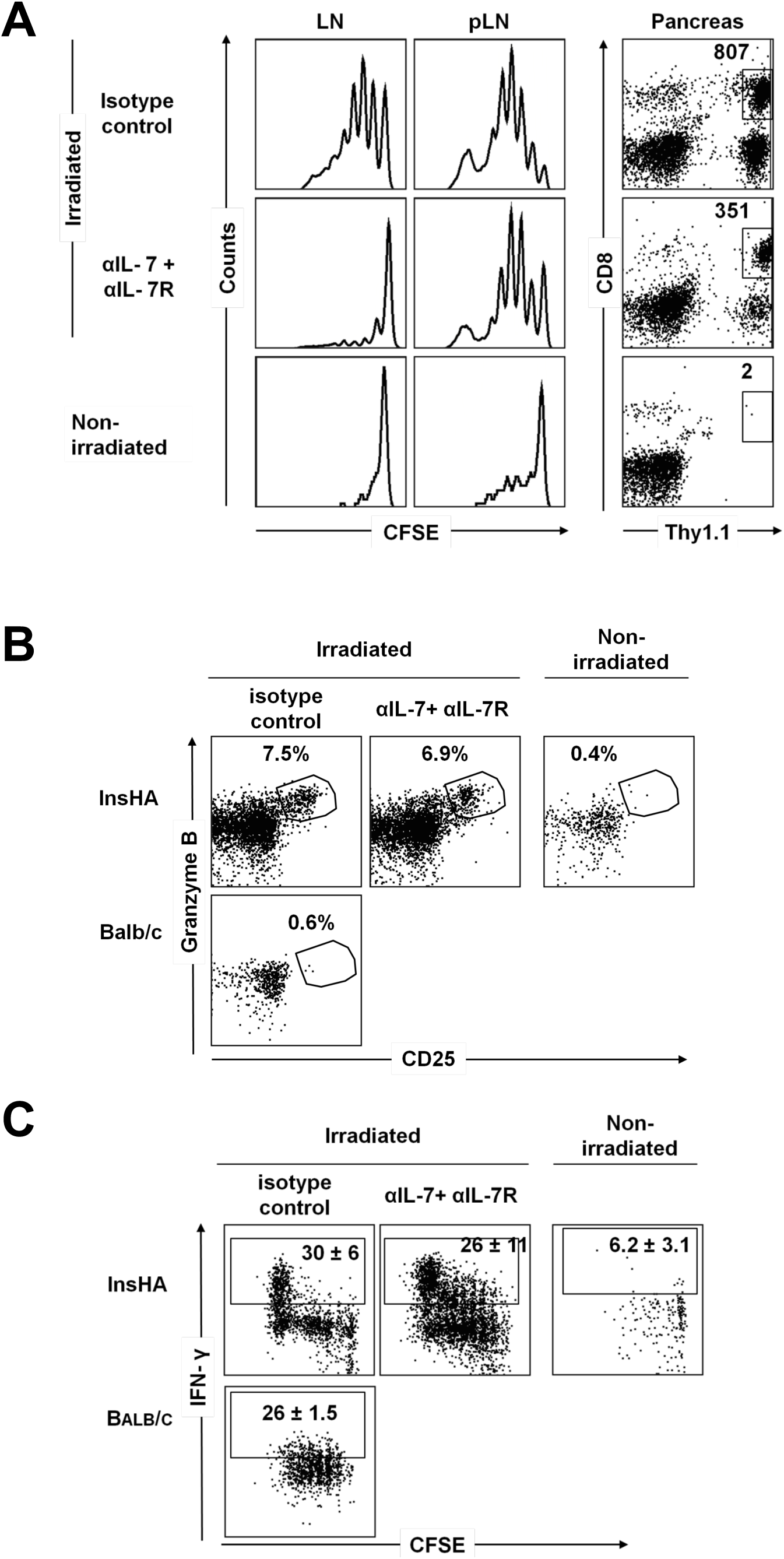
IL-7 blockade does not prevent antigen-driven proliferation and differentiation of Clone 4 CD8^+^ T cells into effectors in irradiated InsHA mice. (**A)** Irradiated InsHA mice adoptively transferred with 5×10^6^ CFSE-labeled naïve Clone 4 Thy1.1^+^ CD8^+^ T cells and 5×10^6^ HNT CD4^+^ T cells were injected with either anti-IL-7 and IL-7Ralpha or isotype control mAbs. Adoptively transferred non-irradiated InsHA served also as controls. Mice were sacrificed on day 8 after transfer and the levels of CFSE fluorescence on gated donor CD8^+^ Thy1.1^+^ lymphocytes in pancreatic lymph nodes (pLN) and lymph nodes (LN) are shown. The presence of CD8^+^ Thy1.1^+^ donor T cells in the pancreas was evaluated by FACS. Numbers indicate FACS event counts in the depicted gates. (**B)** The percentages of Granzyme B^+^ CD25^+^ CD8^+^ donor T cells were analyzed by flow cytometry in pLN at day 8 post transfer. (**C)** On day 8 after transfer, cells from the pLN were incubated with K^d^ HA peptide for 5 h at 37 °C and analyzed by FACS to detect IFN-γ production. Background in non-stimulated controls was less that 3%. The percentages of Clone 4 CD8^+^ T cells producing IFN-γ are indicated as means ± SD (*n* = 3 per group). Data from one representative experiment out of 4 are presented.

**Figure 3.**
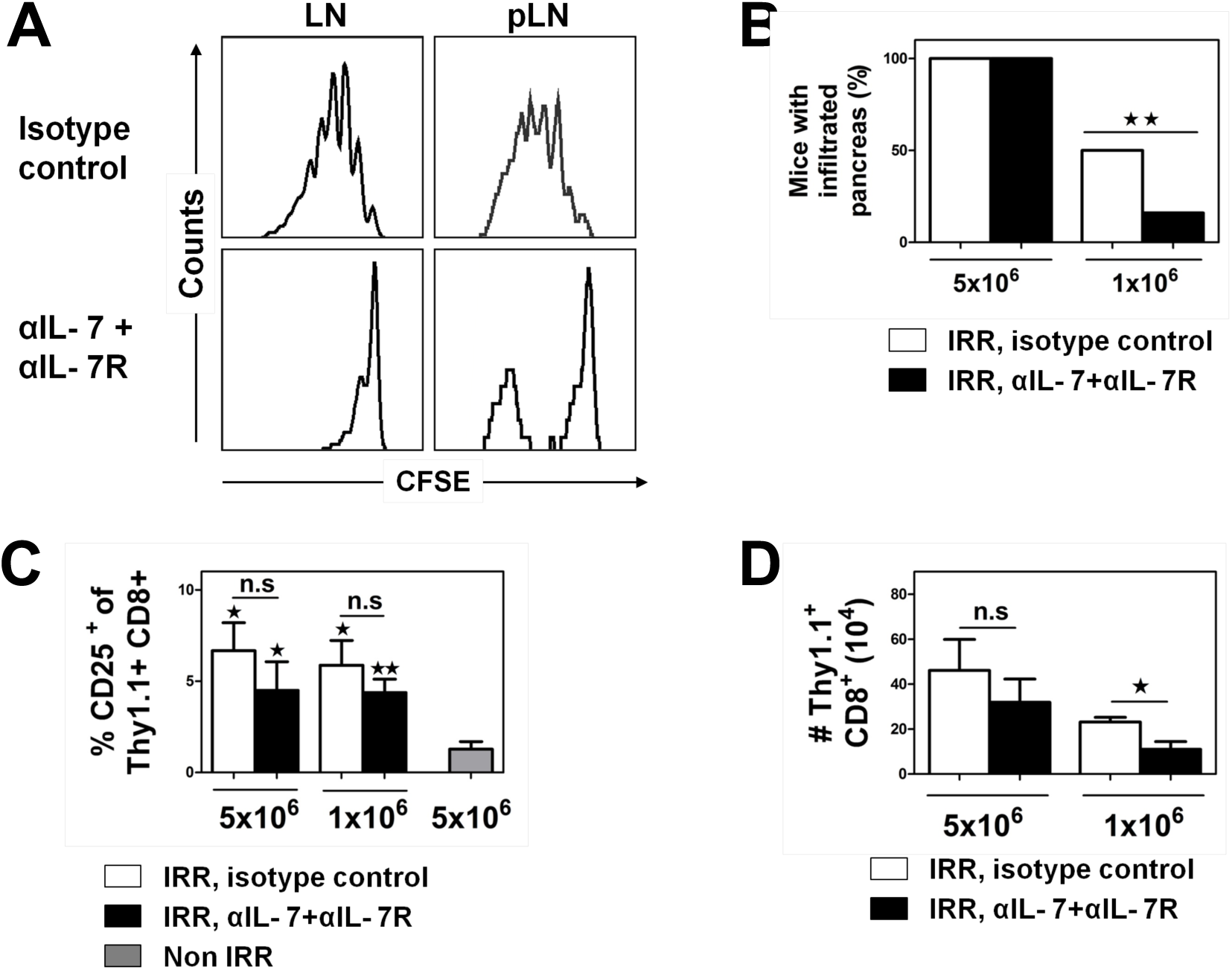
IL-7 blockade does not prevent antigen-driven proliferation and differentiation of Clone 4 CD8^+^ T cells into effectors at low T cell frequencies. (**A)** Irradiated InsHA mice adoptively transferred with 10^6^ CFSE-labeled naïve Clone 4 Thy1.1^+^ CD8^+^ T cells and 10^6^ HNT CD4^+^ T cells were injected with either anti-IL-7 and IL-7Ralpha or isotype control mAbs. Mice were sacrificed at day 8 after transfer and cells from the LN and pLN were analyzed by FACS. Histograms represent CFSE labeling on gated CD8^+^ Thy1.1^+^ lymphocytes. (**B)** Pancreas infiltration by CD8^+^ Thy1.1^+^ T cells was evaluated by FACS at day 8 post-transfer following the injection of the indicated amount of Clone 4 CD8^+^ T cells and HNT CD4^+^ T cells together with isotype control (n=10, cumulative data from 3 independent experiments) (□) or with anti-IL-7 and IL-7Ralpha mAbs (n=10, cumulative data from 3 independent experiments) (▪). A pancreas was considered not to be infiltrated if less than 10 events were detected. The percentages of mice with infiltrated pancreas are shown. (**C)** CD8^+^ Thy1.1^+^ from the pLN of irradiated InsHA mice described in **B** were assessed for expression of CD25 at day 8. Adoptively transferred non-irradiated BALB/c mice (grey bar) served as negative controls. The percentages of CD8^+^ Thy1.1^+^ T cells expressing CD25 are indicated as means ± SD (*n* = 3 per group). (**D)** Total numbers of CD8^+^ donor T cells in the lymphoid organs were enumerated at day 8 post-transfer in mice from groups described in **B**. Results are presented as means ± SD (*n* =3). Data from one representative experiment out of 5 are presented.

Next, we assessed the phenotype of proliferating Clone 4 CD8^+^ T cells in the pLN. CD4 help induced the differentiation of highly proliferating memory-like CD8^+^ T cells into effectors as demonstrated by the upregulation of CD25 and Granzyme B and the dowregulation of CD62L in isotype control mice (Fig. 2B and 3 and data not shown)^21^. Notably, IL-7 blockade did not prevent the differentiation of Clone 4 CD8^+^ T cells in response to HA cross-presentation, since effector cells were generated as efficiently in both groups of mice (Fig. 2B and 3). As expected, Clone 4 CD8^+^ T cells from the pLN of non-irradiated InsHA mice and antigen-free irradiated Balb/c mice^21^ did not differentiate into effector cells (Fig. 2B and 3). Finally, we assessed the ability of Clone 4 donor T cells to produce the effector cytokine IFN-gamma and found no significant differences upon IL-7 blockade (Fig. 2C). It is important to note that Clone 4 cells in non-irradiated InsHA mice do not produce IFN-gamma upon self antigen encounter and that memory-like Clone 4 cells from irradiated BALB/c mice produce this cytokine although the amount produced is lower than in irradiated InsHA mice (Fig. 2C). Taken together, our results show that blockade of Clone 4 CD8^+^ T cell LIP and differentiation into memory-like cells does not prevent the breakdown of cross-tolerance and their differentiation into pathogenic effector CTL in irradiated InsHA mice.

### IL-7 blockade decreases CD8^+^ T cell mediated self-reactivity in irradiated InsHA mice at low T cell frequencies

Our results indicated that differentiation into memory-like cells induced by LIP does not represent an advantage for the breakdown of CD8^+^ T cell cross-tolerance in irradiated mice. However, IL-7 mediated LIP has been shown to enhance CD8^+^ T cell responses^39^. We hypothesized that this could be the result of increasing total numbers of responding T cells. Since the above described experiments were performed transferring relatively high numbers of donor T cells, blocking LIP would not have had any effect on the induction of Clone 4-mediated self-reactivity. Therefore, we next performed similar experiments utilizing lower numbers of donor T cells. Irradiated InsHA mice were injected with 10^6^ of each Clone 4 CD8^+^ and HNT CD4^+^ T cells and were treated with M25 and A7R34 or isotype control mAbs. In this case, only 71% of the mice in the control group developed disease, showing that the amount of transferred T cells is close to the threshold required to induce disease (Table 1). In these conditions, IL-7 blockade reduced disease incidence down to 38% of treated mice (Table 1). In agreement with these results, we found that IL-7 blockade reduced by 40% the number of mice in which donor CD8^+^ T cells were detected infiltrating the pancreas at day 8 post transfer (Fig. 3B).

In order to shed light in the differences on self-reactivity observed at low doses of donor T cells, we next analyzed proliferation, activation status and numbers of clone 4 CD8^+^ T cells in lymphoid organs. We found that IL-7 blockade prevented LIP in non-draining lymph nodes. However, as observed at high donor T cell numbers, treatment did not prevent the antigen-induced fast proliferation in the pLN (Fig. 3A). Furthermore, Clone 4 CD8^+^ T effector cells were generated in the pLN of all mice analyzed, as evidenced by the generation of a CD25^+^ subpopulation, and no significant differences in the rate of differentiation from treated and non-treated animals was observed (Fig. 3C). However, treatment reduced significantly by 3-fold the total numbers of donor CD8^+^ T cells recovered from the LN (Fig. 3D). Collectively, our data suggest that at low T cell frequencies, blocking LIP decreased self-reactivity by reducing the availability of potentially auto-reactive CD8^+^ T cells without preventing the breakdown of self-tolerance and their ability to differentiate into effector cells.

### IL-7 blockade does not release regulatory mechanisms that prevent self-reactivity in irradiated InsHA mice

It has been recently shown that IL-7 blockade may result in the induction of regulatory mechanisms and prevent T cell mediated autoimmunity^38,40^. To assess whether this phenomenon contribute to the decreased self-reactivity observed in our model at low T cell frequencies, we examined levels of Tregs in treated versus non-treated InsHA mice. Interestingly, we found that IL-7 blockade promoted a moderate relative increase of endogenous CD4^+^ FoxP3^+^ regulatory T cells in InsHA mice (Fig. 4). An increase in Tregs could contribute, together with the decrease in the total numbers of potentially autoreactive T cells observed in treated mice, to reduce the onset of autoimmune diabetes in irradiated InsHA mice. To test this hypothesis, we transferred polyclonal, syngeneic Tregs into irradiated InsHA injected with 10^6^ Clone 4 CD8^+^ and HNT CD4^+^ T cells. We found that this transfer did not reduce self-reactivity compared to control mice (Table 2). As a control and to verify the efficacy of our isolated Tregs, we transferred them into antibody-T cell depleted InsHA mice injected with Clone 4 CD8^+^ and HNT CD4^+^ T cells. Under these conditions, InsHA mice develop autoimmunity as in irradiated mice (our unpublished results). In this setting, polyclonal, syngeneic Tregs were able to completely prevent the onset of autoimmune diabetes (Supplemental Table 1). These results demonstrate that increasing Treg numbers in irradiated InsHA mice does not reduce T cell mediated self-reactivity.

**Figure 4.**
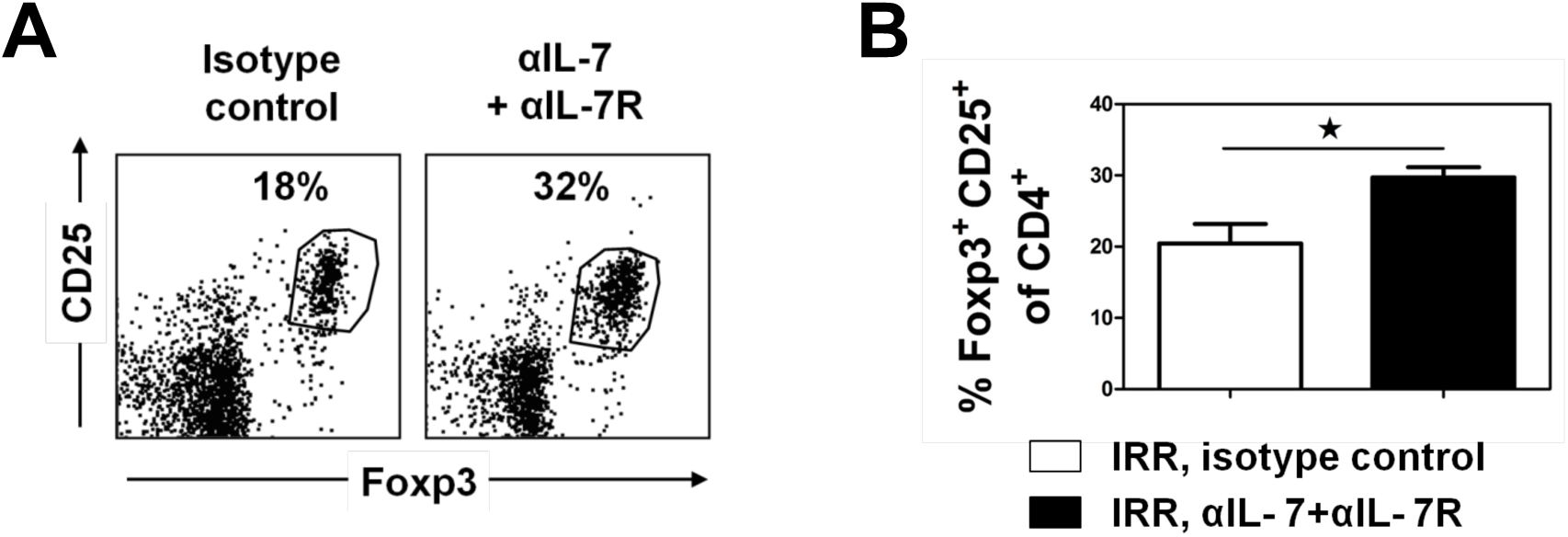
IL-7 blockade promotes enhances the relative numbers of Tregs in irradiated InsHA mice. Mice from groups described in Figure 3A were sacrificed at day 8 after transfer and expression on CD25 and Foxp3 was assessed on endogenous CD4^+^ T cells in the LN. **(A)** Plots of representative mice from each group indicating the percetages of CD4^+^ CD25^+^ Foxp3^+^ T cells. **(B)** Data represents the mean percentages ± SD (*n* =3). Data from one representative experiment out of 2 are presented.

**Table 2.**
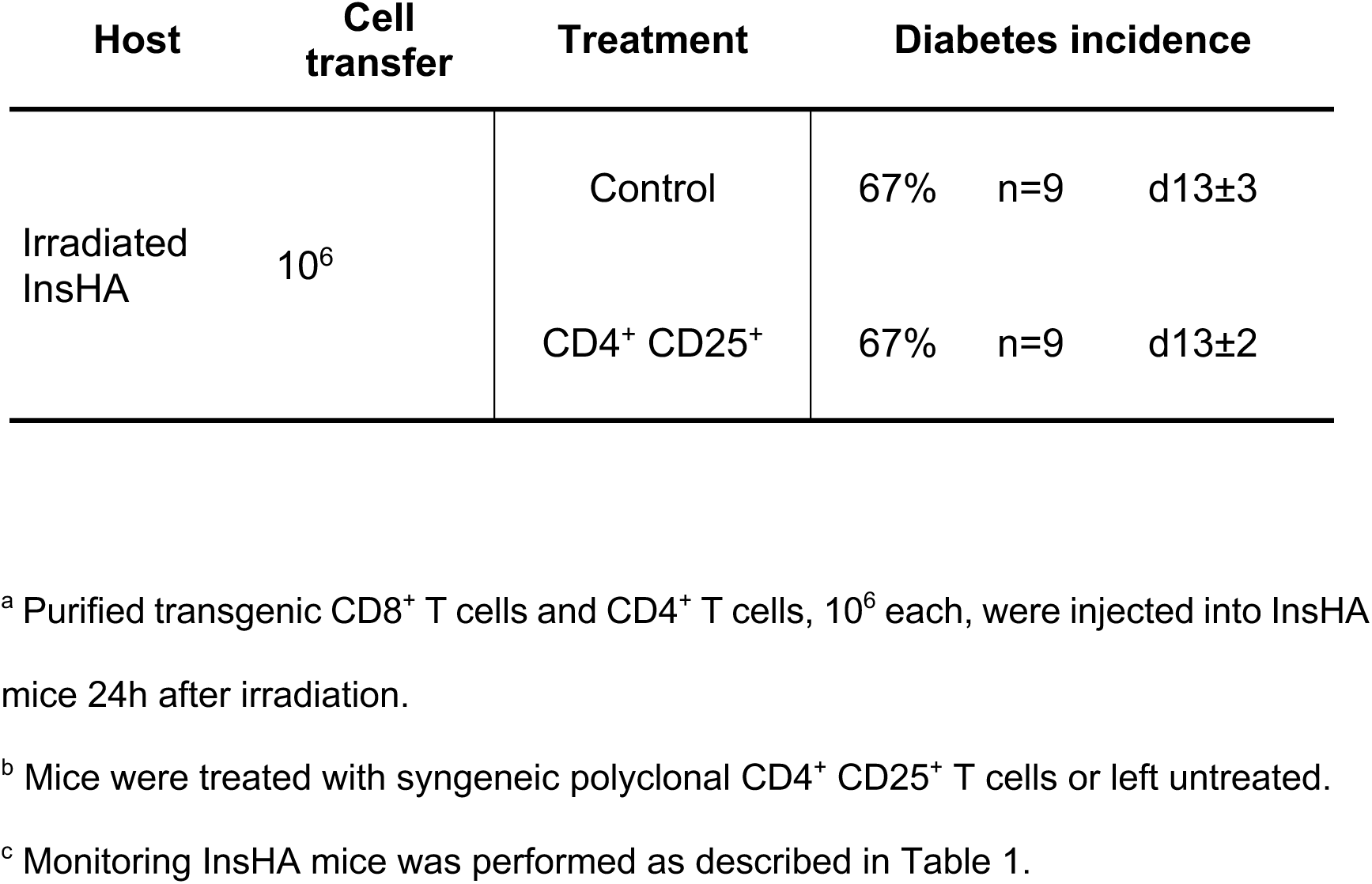
Tregs do not prevent self-reactivity in irradiated InsHA mice.

| Host | Cell transfer | Treatment | Diabetes incidence |  |  |
| --- | --- | --- | --- | --- | --- |
| Irradiated InsHA | 10<sup>6</sup> | Control | 67% | n=9 | d13±3 |
|  |  | CD4<sup>+</sup> CD25<sup>+</sup> | 67% | n=9 | d13±2 |

<sup>a</sup> Purified transgenic CD8<sup>+</sup> T cells and CD4<sup>+</sup> T cells, 10<sup>6</sup> each, were injected into InsHA mice 24h after irradiation.
<sup>b</sup> Mice were treated with syngeneic polyclonal CD4<sup>+</sup> CD25<sup>+</sup> T cells or left untreated.
<sup>c</sup> Monitoring InsHA mice was performed as described in Table 1.

### Enhanced LIP augments CD8^+^ T cell mediated self-reactivity in irradiated InsHA mice

To determine whether an increase in donor T cell numbers through LIP augments self-reactivity in irradiated InsHA mice, we sought to potentiate LIP of Clone 4 CD8^+^ T cells by providing exogenous IL-7. Given the short half-life of recombinant IL-7 *in vivo*, we utilized an IL-7/anti-IL-7 mAb immune complex (IC) for the treatment of mice. It has been shown that IC enhances the potency of IL-7 mainly by increasing its half-life^33,41^. Indeed, provision of the IC to irradiated BALB/c mice adoptively transferred with naïve Clone 4 CD8^+^ T cells enhanced LIP without altering their differentiation into memory-like cells (Sup. Fig. 2). Next, we assessed the effect of IL-7 IC treatment in the onset of autoimmune diabetes in irradiated InsHA mice injected with 0.5×10^6^ of each naive Clone 4 CD8^+^ and CD4^+^ T cells. With these low numbers of transferred HA-specific T cells only 57% of untreated control mice developed disease. Notably, IL-7 IC treatment augmented up to 85% the onset of disease in irradiated InsHA mice (Table 3). This was associated with an enhanced accumulation of Clone 4 CD8^+^ T cells in the LN (Fig. 5A). Importantly, IL-7 IC did not significantly change the rate of differentiation of Clone 4 CD8^+^ T cells into effectors in the pLN (Fig. 5B). Thus, IL-7 enhances LIP in irradiated InsHA mice promoting an increase of potentially autoreactive CD8^+^ T cell numbers that, in turn, correlates with an augmentation in self-reactivity.

**Figure 5.**
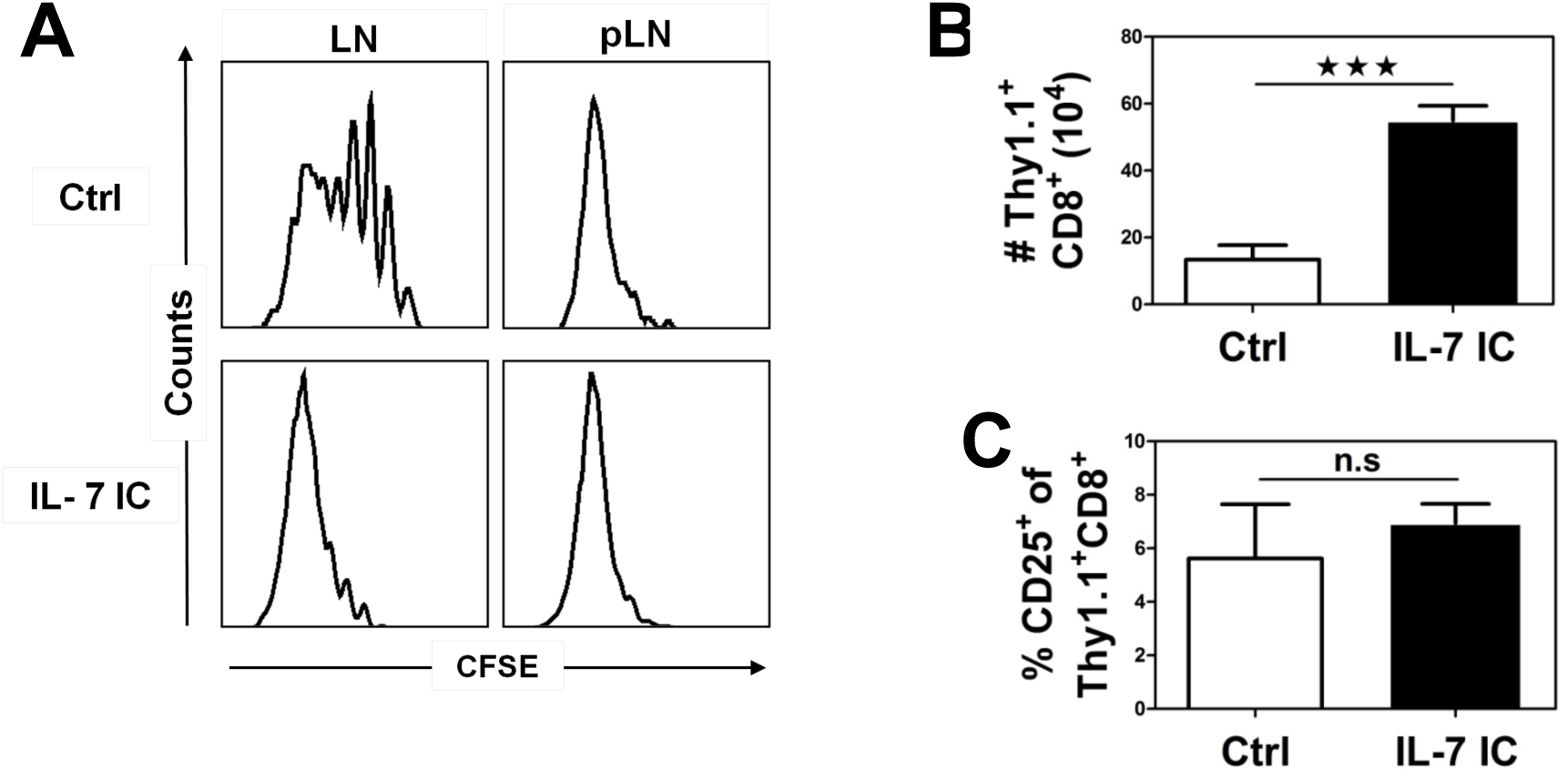
IL-7 immune complex enhances Clone 4 CD8^+^ T cells numbers in irradiated InsHA mice. **(A)** Irradiated InsHA mice injected with 0.5×10^6^ CFSE-labeled naïve Clone 4 Thy1.1^+^ CD8^+^ and 0.5×10^6^ HNT CD4^+^ T cells and were treated with IL-7/M25 anti-IL-7 immune complex (I.C.) or left untreated. Mice were sacrificed on day 8 after transfer. Levels of CFSE fluorescence on gated CD8^+^ Thy1.1^+^ donor lymphocytes in LN and pLN are shown. **(B)** Absolute numbers of CD8^+^ donor T cells in the lymphoid organs were enumerated in mice treated with with IL-7 I.C. (▪) and control mice (□). Results are presented as means ± SD (*n* = 4-8). **(C)** Plots represent expression of CD25 on gated CD8^+^ Thy1.1^+^ lymphocytes in the pLN (n=4-5). Data from 3 independent experiments are presented.

**Table 3.** IL-7 I.C. enhances self-reactivity in InsHA mice.

| Host | Cell transfer<sup>a</sup> | Treatment<sup>b</sup> | Diabetes incidence<sup>c</sup> |  |  |
| --- | --- | --- | --- | --- | --- |
| Irradiated<br>InsHA | 0,5.10<sup>6</sup> | Control | 57% | n=14 | d11±3 |
|  |  | I.C IL- 7 | 85% | n=14 | d11±2 |

<sup>a</sup> Purified transgenic CD8<sup>+</sup> T cells and CD4<sup>+</sup> T cells, 0.5x10<sup>6</sup> each, were injected into InsHA mice 24h after irradiation.
<sup>b</sup> Mice were treated with IL-7 IC or left untreated.
<sup>c</sup> Monitoring InsHA mice was performed as described in Table 1.

## DISCUSSION

Currently, irradiation and lymphodepleting drugs are utilized as conditioning protocols to enhance T cell based immunotherapies in cancer patients, although the basis underlying their beneficial effects are not well understood. The fact that many tumor-asociated antigens are proteins also expressed in some normal tissues and that lymphopenia is a feature of many models of autoimmunity pointed to lymphopenia-inducing protocols as perturbators of the mechanisms of peripheral T cell tolerance^22,23^. The environment generated after irradiation in the immune system supports extensive T cell LIP and differentiation into memory-like cells^34^. Thus, it was hypothesized that LIP, by driving CD8^+^ T cells into a more activated state, could induce a breakdown of self-tolerance and explain the enhanced CTL responses observed^22,23^. However, the fact that lymphopenia-inducing protocols, such as irradiation, have multiple effects on the organism have made difficult to assess separately the role of LIP. Brown et al. reported that in RAG deficient mice, in which LIP is thought to be uncoupled from other effects induced by irradiation, soluble peptide-induced CD8^+^ T cell anergy could be reversed^42^. In contrast, it has been recently shown that while irradiation could revert the anergic state of CD8^+^ T cells induced by self-antigen in the neonatal period, LIP in RAG deficient mice is not sufficient to do so^5^. This discrepancy was explained by the fact that self-antigen induces a deeper anergic state than soluble peptide^5^. Our results show for the first time that LIP in irradiated mice is not required to overcome deletional tolerance induced by self-antigen cross-presentation. Potentially autoreactive CD8^+^ T cells that remained in a naïve state, upon IL-7 blockade, responded to antigen and CD4 help differentiating into effector CTL in the pLN as efficiently as memory-like cells. Our data and that from Oelert et al. clearly indicate that other signals generated after irradiation are responsible for the breakdown of tolerance induced by self-antigens. Likely candidates are the systemic activation of antigen presenting cells induced by commensal bacteria translocation^43^ or the increased availability of cytokines other than IL-7 observed in irradiated mice^5,44^. Future studies will help define whether these or additional yet not identified signals are responsible for the breakdown of CD8^+^ T cell tolerance in irradiated mice.

The main roles of IL-7 are arguably mediating lymphocyte development and the maintenance of T cell homeostasis^39^. An additional new role as an enhancer of anti-viral, -tumor and -self T cell responses is emerging for this cytokine from recent studies in models in which its levels or signaling were augmented^20,45–49^. To the known capacity of increasing T cell numbers and therefore the magnitude of T cell responses, an increased IL-7 availability can counteract regulatory mechanisms that otherwise control T cell effector functions^38,40,45,46^. IL-7 has been shown to promote Th17 and Th1 differentiation, which can contribute to the pathogenesis of certain autoimmune diseases^40,45^. Of interest, Lee et al. showed that IL-7 can promote the in vitro differentiation of NOD CD8^+^ T cells into effectors upon agonist TCR engagement^40^. Thus, multiple mechanisms can account for the enhanced T cell responses observed when increased levels of IL-7 are present. The data presented here demonstrate that in the presence or absence of IL-7 signaling the rate of differentiation of Clone 4 CD8^+^ T cells into CTL after antigen encounter in the pLN is similar. This suggests that the levels of IL-7 present in irradiated mice do not support a qualitative gain of effector functions. This is in agreement with previous studies in which exogenous IL-7 did not significantly enhance Clone 4 CD8^+^ T cell functionality in non irradiated mice^50^. It is interesting to speculate that the enhanced effector functions observed with CD8^+^ T cells from NOD mice in the presence of IL-7 could be related to the presence of polymorphisms that could modify IL-7 signaling.

It has been recently shown that IL-7 blockade prevents disease progression in the NOD spontaneous model of autoimmune diabetes^38,40^. Long term IL-7 signaling blockade induced upregulation of the negative regulator PD-1 on CD4^+^ Th effector cells that dampened the Th1 pathogenic response^38,40^. Treatment also resulted in increased levels of CD4^+^ FoxP3^+^ regulatory cells that in turn may contribute to counterbalance effector cells and revert disease progression^38,40^. Increased levels of Tregs in treated mice were likely the reflection of their lower expression of IL-7R and less dependence on IL-7 signals for survival compared to conventional T cells. In the NOD model, CD4^+^ T cells have been described to be pathogenic in themselves and PD-1 upregulation was not assessed on CD8^+^ T cells in those reports^48,50^. In our model only CD8^+^ T cells are pathogenic effectors. CD8^+^ T cells are also susceptible to regulation by engagement of PD-1. However, we did not see a differential upregulation of PD-1 on Clone 4 CD8^+^ T cells upon IL-7 signaling blockade. Whether these differences are T cell subset related or dependent on the self-reactivity predisposing genetic background in NOD mice deserves further attention. Notably, we did observe a relative increase in the levels of Tregs in treated mice. Nonetheless, transfer of high numbers of Tregs to non-treated mice did not reduce self-reactivity, indicating that modest variations in their levels are not responsible for the differences in self-reactivity observed in irradiated mice. Our results corroborate previous studies demonstrating that drastic variations in the levels of Tregs are needed to observe a loss of self-tolerance in adult mice^51,52^.

In conclusion, our results demonstrate that CD8^+^ T cell LIP is not required to overcome deletional tolerance induced by self-antigen cross-presentation in irradiated mice. CD8^+^ T cells that do not differentiate into memory-like cells are able to become effector CTL due to additional cues provided by irradiation. However, LIP plays an important role in enhancing self-reactivity through increasing the frequency of potentially autoreactive T cells. Our results are consistent with and could help explain recent reports showing that i) anti-tumor responses could be enhanced by LIP, ii) the onset of autoimmune diabetes mediated by endogenous CD8^+^ T cells can be induced by increasing the frequency of self-antigen-specific endogenous CD8^+^ T cells though LIP and iii) human autoimmunity in lymphopenic patients correlates with LIP^5,20,53^. These data have important implications for the understanding of autoimmune processes under lymphopenic conditions as well as for the development of cancer immunotherapies.

## ACKNOWLEDGMENTS

We are indebted to the MRI-RIO imaging platform (GIS-IBISA, Languedoc-Roussillon) for flow cytometry experiments and the T&TA core facilities for animal experiments. This work was supported by grants from the European Community SUDOE-FEDER (Contract # IMMUNONET SOE1/P1/E014) and La Ligue Contre le Cancer Hérault to JH.

## Author contributions

Designed experiments: JH. Performed the experiments: MV, CLS, GEC. Analyzed the data: MV, CLS, GEC, JH. Wrote the paper: JH.

