## Supplemental figures and tables for "Differentiation of naïve into memory-phenotype CD8^+^ T cells does not promote the breakdown of peripheral tolerance in irradiated mice"

**Supplemental Table 1. Tregs prevent self-reactivity in T cell depleted InsHA mice.**

| Host[a] | Cell transfer[b] | Treatment[c] | Diabetes incidence[d] | | |
|---|---|---|---|---|---|
| Anti-Thy1.2 InsHA | $5x10^6$ | Ctrl | **100%** | n=5 | d9±1 |
| | | $CD4^+CD25^+$ | **0%** | n=5 | |

[a] InsHA mice were injected i.p. with 0.5 mg per mouse per dose of anti-Thy1.2 mAb (clone 30H12, BioXcell) on days 8 and 4 before adoptive T cell transfer.

[b] Equal numbers of purified transgenic Clone 4 $CD8^+$ and HNT $CD4^+$ T cells were injected as indicated into T cell depleted InsHA mice.

[c] Mice were injected with $1.5x10^6$ polyclonal syngeneic $Thy1.1^+$ $CD4^+$ $CD25^+$ T cells 12h after they received the mAb or were left untreated.

[d] The onset of autoimmunity was evaluated by measuring blood glucose levels. Mice were followed over a 30 day period and were considered diabetic when levels were above 300 mg/dl in two consecutive measurements. The day of disease onset (d) is indicated when applicable. Data from two independent experiments is presented.

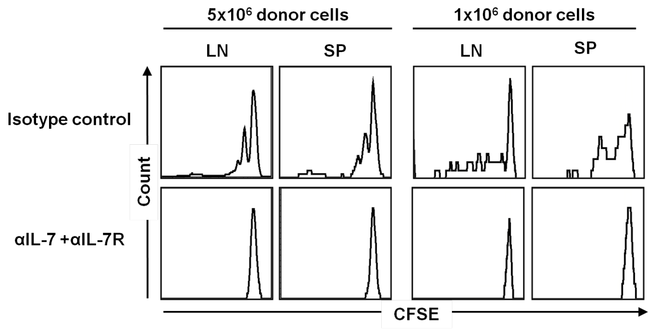


**Supplemental Figure 1. IL-7 blockade prevents LIP of HNT CD4$^+$ T cells.** (**A**) Irradiated BALB/c mice were injected with equal numbers of CFSE-labeled naïve Clone 4 Thy1.1$^+$ CD8$^+$ T cells and HNT CD4$^+$ T cells as indicated and treated with either anti-IL-7 and IL-7Ralpha or isotype control mAbs. Mice were sacrificed at day 8 after transfer and cells from the LN and spleen were analyzed by FACS. Histograms represent CFSE labeling on gated CD4$^+$ Thy1.1$^+$ lymphocytes. Data from one representative experiment out of 3 are depicted.

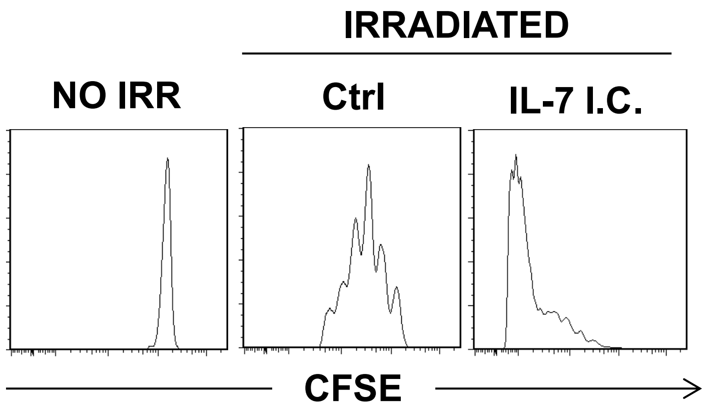


**Supplemental Figure 2. IL-7 immune complex enhances LIP of Clone 4 CD8$^{+}$ T cells.** Irradiated BALB/c mice injected with 0.5x10$^{6}$ CFSE-labeled naïve Clone 4 Thy1.1$^{+}$ CD8$^{+}$ and 0.5x10$^{6}$ HNT CD4$^{+}$ T cells were treated with IL-7/M25 anti-IL-7 immune complex (I.C.) or left untreated. Mice were sacrificed on day 8 after transfer. Levels of CFSE fluorescence on gated CD8$^{+}$ Thy1.1$^{+}$ donor lymphocytes in LN are shown. Data from one representative experiment out of two are depicted.
